# A Two-State Epistasis Model Reduces Missing Heritability of Complex Traits

**DOI:** 10.1101/017491

**Authors:** Kerry L. Bubb, Christine Queitsch

## Abstract

Despite decade-long efforts, the genetic underpinnings of many complex traits and diseases remain largely elusive. It is increasingly recognized that a purely additive model, upon which most genome-wide association studies (GWAS) rely, is insufficient. Although thousands of significant trait-associated loci have been identified, purely additive models leave much of the inferred genetic variance unexplained. Several factors have been invoked to explain the ‘missing heritability’, including epistasis. Accounting for all possible epistatic interactions is computationally complex and requires very large samples. Here, we propose a simple two-state epistasis model, in which individuals show either high or low variant penetrance with respect to a certain trait. The use of this model increases the power to detect additive trait-associated loci. We show that this model is consistent with current GWAS results and improves fit with heritability observations based on twin studies. We suggest that accounting for variant penetrance will significantly increase our power to identify underlying additive loci.

## Introduction

Like Mendelian traits, many complex traits, including autism, schizophrenia and cleft lip/palate, are often present in one of two states -- affected or unaffected. Unlike Mendelian traits, for which the underlying genetics is binary, myriad genetic and environmental factors may determine complex trait status. Such complex traits that rely on thresholding of an underlying, often hidden liability, are commonly called threshold characters [1–3], or threshold traits.

GWAS is the most common approach to discover the multiple loci underlying threshold traits, relying on the assumption that risk alleles are more frequent among affected individuals than unaffected individuals. As currently implemented, GWAS also assume that loci contribute independently and additively to a complex trait. How successful this experimental design has been is a matter of debate. Since 2007, GWAS have identified thousands of loci with significant trait or disease association and implicated previously unconnected pathways in disease processes [4]. Examples of clear biological and clinical relevance include the autophagy pathway in Crohn’s disease [5–8] and the JAK-STAT signaling pathway in rheumatoid arthritis [9,10], among others. However, most trait-associated loci individually explain very little of the inferred genetic variance. They are therefore of limited use for predicting the disease risk of a given individual and for understanding the mechanistic underpinnings of complex traits. It is widely acknowledged that there is room for improvement [11,12].

Many hypotheses have emerged to explain the ‘missing heritability’ [13–15]. Some complex diseases such as autism spectrum disorder represent likely insufficiently resolved pools of phenotypically similar, but inherently rarer, disease traits with different genetic underpinnings. If so, the observed odds ratios for significantly trait-associated SNP are low because a common, trait-associated SNP is linked to a rare causative mutation that only appears in a small subset of haplotypes [16]. Fine-grained phenotyping rather than relying on discrete, binary diagnoses should help to explore this hypothesis [17]. Some have suggested that the heritability of complex traits is overestimated [18,19]. Studies accounting for all SNPs genome-wide simultaneously, as opposed to individually associating SNPs with traits, indicate that this explanation is unlikely for many traits [20,21]. Others have invoked currently inaccessible genetic or structural variants or rare risk alleles of moderate effect as major factors in complex traits [13,14,22]. However, at least for autism, recent studies suggest that common variants account for over 50% of the genetic risk [23,24].

Finally, although additive genetics is certainly a major fodder for evolution and selective breeding since epistatic interactions often present as additive [25,26], epistasis can have large influence on complex traits [27]. An excellent example of the importance of epistasis comes from plant breeding. As breeders increased seed yield, presumably via additive genetic factors, seeds became far more numerous, larger, and heavier. The increasing pressure on plant stalks required new mutations that enabled plants to remain erect under the increased seed weight -- this epistatic interaction enabled the Green Revolution [28] that vastly increased food security in many poor parts of the world [29].

In humans, twin concordance rates indicate that at least some of the genetic variation influencing complex diseases is non-additive. Experimental evidence demonstrates that epistasis, *i.e*. the phenotypically relevant, and often non-reciprocal interaction of non-allelic genes, is pervasive in complex traits in various model organisms [30–32]. There is no reason to assume that the genetic architecture of complex traits differs between humans and other highly complex eukaryotes. Therefore, the inclusion of epistatic effects in statistical models has been increasingly suggested and even attempted in some studies [27,33,34]. Although models that allow for all gene × gene (and gene × gene × gene, etc) interactions will be more realistic, such a higher-order models require much larger datasets and faster algorithms [33,35–39].

Studies in model organisms suggest that a simpler, two-state epistasis may apply to complex traits and diseases [32,40]. In contrast to the familiar often small-effect, gene x gene interactions, certain genetic and environmental factors can act as strong modifiers for many other loci [30,41–46]. In addition, the activity of strong genetic modifiers can be modulated by environmental stress [47]. Based on studies in plants, worms, and yeast, the number of strong genetic modifiers is small, possibly ~10% of all genes [30,46,48].

Although the existence of differences in variant penetrance among individuals is not a new concept, its effect on GWAS in humans has not been investigated. Here, we present a simple two-state epistasis model, in which binary disease status of an individual depends on a combination of additive alleles (as before) as well as their penetrance in a given individual. Such a model might be called an A×P, or Additive × Penetrance model. A population consisting of all individuals with increased penetrance of many different genetic variants will have higher phenotypic variation, *i.e*. it will be less phenotypically robust, than an otherwise equivalent population consisting of all robust individuals with low penetrance [31]. Theoretical population genetics provides a strong argument for the potential benefits of maintaining a population with a balance of robust and non-robust individuals [49–55]. As we show, this model increases the power to detect additive trait-associated loci and improves fits with heritability observations based on twin studies. Although it is perhaps unsurprising that adding a parameter to a model improves its predictive power, that is precisely our point --only *one* parameter, not *g^2^*, *g^3^*,… interaction parameters (where *g* is the number of genes in the genome), results in a marked improvement of fit.

In the absence of robustness measures in humans, one may argue that such model is of limited use to improve GWAS in humans. However, we argue that our model’s success should inform our approach to finding disease-causing loci and we discuss strategies how to apply it to existing and future GWAS data.

## Results

**Heritability estimates imply substantial non-additivity of genetic factors contributing to human disorders:** For a quantitative trait, the fraction of phenotypic variance due to *additive* genetic factors (narrow-sense heritability, or *h^2^*) is straightforward to calculate: it is a function of the slope of the line of regression between pairs of related individuals (**Fig.1A**) [3,56]. The fraction of phenotypic variance due to *all* genetic factors (broad-sense heritability, or *H^2^*) is less straightforward to calculate (see Supplement) [19,36].

**Figure 1:**
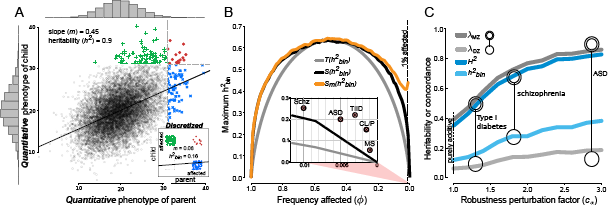
**Comparison of *narrow-sense* heritability of quantitative liability, *observable* heritability of discrete traits, and *empirically determined* heritability of certain complex threshold traits.** (**A**) Each symbol in the large scatter plot represents the genetic liabilities for one parent-child pair. Each symbol in the inset scatter plot represents the disease state for one parent-child pair (disease state must be 0=unaffected or 1=affected; points were jittered for visual clarity). Green pluses indicate cases in which the child is affected (*i.e*. has crossed the diagnostic threshold) and the parent is not affected; blue crosses indicate cases in which the parent is affected and the child is not; red filled circles indicate cases in which both are affected; black open circles indicate cases in which neither is affected. Input parameter values for the simulation were as follows: number of additive risk-loci (*n*) = 100; frequency of additive risk-allele (*p*) = 0.1; frequency of non-robust state (*p_r_*) = 0. (**B**) The relationship between frequency of affected individuals in the population (φ) and the maximum possible observable *binary* heritability (*h*^2^_*bin*_) when the underlying liability is entirely determined by additive genetic factors (*h*^2^ = 1) as predicted by theory [57], *T(h^2^_bin_)*, as determined by simulation, *S(h^2^_bin_)*, or as determined by simulation of a population that includes two types of individuals, robust and non-robust, at a ratio of 99:1, *S_m_*(*h*^2^_*bin*_), where *H*^2^ = 1. In the inset, we display the relevant φ range where 1% or less of the population is affected, which is common for many complex diseases. For the diseases shown, empirically determined heritability *O(h^2^_bin_)* is up to twice as high as predicted under the models that do not include an epistatic robustness factor. (**C**) In a population containing just 1% non-robust individuals, increasing the robustness perturbation factor (*c*_α_), *i.e*. the fold-change of the additive genetic liability, increases both heritability and twin concordance to levels observed in several complex diseases.

Further complications arise for traits that are observably discrete, but rely on thresholding of an underlying, unobserved quantitative *liability*. Such traits are called *threshold characters* [1–3]. Using the same method to measure additive heritability as before – doubling the slope of the line of regression between parent-offspring pairs – results in a substantially smaller estimate of the additive heritability (**Fig. 1A, inset**).

The degree to which this *observable* discrete additive heritability (*h*^2^_*o*_) is decreased relative to the heritability of the *underlying* additive liability (*h*^2^) strongly depends on where the diagnostic threshold is drawn – in other words, the fraction of individuals that are affected (*φ*). Henceforth, we use *h*^2^_*bin*_ to refer to the observable heritability of threshold traits (binary, affected/unaffected).

In 1950, Dempster and Lerner [57] derived a formula relating *h*^2^ to *h*^2^_*bin*_ with the simplifying assumption that the quantitative trait (liability) is normally distributed in the population (see *Appendix II*). Here, we refer to this Dempster-Lerner derivation as *T*(*h*^2^_*bin*_) as the maximum heritability of the binary character attainable if the underlying liability consisted purely of additive genetic factors (*h*^2^=1) (**Fig. 1B**).

Expected values for *h*^2^_*bin*_ can also be determined via simulation, in which the additive liability is binomially distributed (*N* is twice the number of additive loci; *P* is the frequency of the risk allele at each locus). We refer to the maximum binary heritability attained via simulation of a *purely additive* model with *h^2^=1* as *S(h^2^_bin_)*.

In either case, binary heritability *h*^2^_*bin*_ reaches a maximum (with respect to *h*^2^) when *half of the population is affected* and drops off rapidly as the fraction of affected individuals approaches low values typical for common discrete complex traits, such as autism, schizophrenia, and multiple sclerosis (**Fig. 1B**). However, *empirically determined h*^2^_*bin*_ values for these traits – *O*(*h*^2^_*bin*_) – are much higher (**Table S1**), calling a purely additive model into question.

To explore the implied non-additive factors, we developed a simulator that designates a subset of the population as non-robust by incorporating a robustness perturbation factor that increases the effect size of all additive risk alleles, *i.e*. increases variant penetrance. Using this simulator, a purely robust population maintains the theoretical relationship between the fraction of affected individuals (*φ*) and *h*^2^_*bin*_ [here called *S*(*h*^2^_*bin*_)] (**Fig. 1B**, black curve). Simulation of a mixed population that includes a subpopulation of non-robust individuals with increased variant penetrance results in an increase of *h*^2^_*bin*_ [here called *S_m_*(*h*^2^_*bin*_)] even when the binary trait is at very low frequency (**Fig. 1B**, yellow curve). Below, we explore the effect of varying this robustness factor, as well as the frequency of non-robust individuals in the population and the contribution of non-genetic factors (*H^2^* < 1).

Hill and colleagues [58] proposed a simpler function to determine the extent of non-additive genetic effects: the correlation of monozygotic twins minus twice the correlation of dizygotic twins (*r_MZ_*-2*r_DZ_*) to evaluate both continuous quantitative traits such as height and threshold traits such as endometriosis. If *r_MZ_>r_DZ_*, resemblance is partly due to genetic factors; *r_MZ_>*2*r_DZ_* implicates non-additive genetic effects. This metric makes intuitive sense, because monozygotic twins share 100% their alleles, and dizygotic twins share 50% of their alleles. While most of the complex traits that Hill and colleagues examine showed *r_MZ_*-2*r_DZ_* values close to or less than zero, for many important threshold traits *r_MZ_*-2*r_DZ_* was greater than zero (**Table S1**).

Therefore, we quantify the non-additive genetic component of complex traits and diseases by comparing *empirically determined* heritability, *O(h^2^_bin_)*, to binary heritability with and without including a robustness perturbation factor [*S(h^2^_bin_)*, additive model, assuming all individuals in the population are equally robust; *S_m_(h^2^_bin_)*, a two-state model, assuming that the population includes a subset of individuals with decreased robustness]. Simulated populations that are a mixture of robust and non-robust individuals produce levels of *h^2^_bin_* that are more consistent with empirically determined *O(h^2^_bin_)* from twin studies (**Fig. 1C, Table S1**).

**Robustness as a two-state model of epistasis:** We implemented a liability model such that each individual in a population has a positive liability that is simply a linear combination of *n* additive genetic factors and noise (ε) (**Fig. 2A**). Individuals with liability in the top fraction of the population (threshold determined by observed incidence levels, *φ*) are assigned affected status.

**Figure 2:**
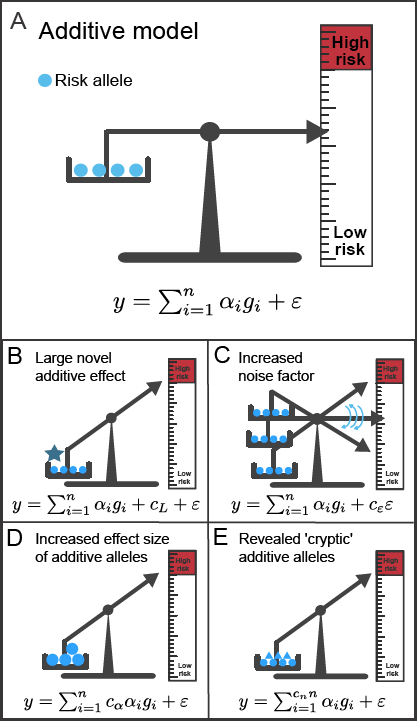
**Base additive model and modifications.** (**A**) Base additive model. (**B**) Additive model plus single large-effect factor (*c_L_*). (**C**) Additive model with increased noise, *i.e*. increased residual effect (*c_ε_*). (**D**) Additive model assuming existence of non-robust state that increases effect size of additive risk alleles (*c_α_*). (**E**) Additive model assuming existence of non-robust state that reveals previously phenotypically silent (cryptic) risk alleles (*c_n_*), represented by the additional blue triangles. [Note: *y* is some quantitative phenotype, *α_i_*’s are the weights of each of the contributing genetic components, *n* is the number of contributing genetic components, *c* is the residual effect, which is assumed to be normally distributed with mean zero and standard deviation *ρ_E_*, and *g_i_* is an indicator variable, taking values 0 or 1 depending on whether the risk allele is present at the *i*-th locus.]

We then generated mixed populations in which some fraction of the population is designated as non-robust and modify their liability in one of four ways, each increasing the fraction of affected individuals that are non-robust (see **Supplementary Methods** for details). First, we assigned a novel liability factor of large additive effect to our non-robust individuals (**Fig 2B**). Second, we increased the fraction of liability due to noise (**Fig 2C**). Third, we increased the effect size of additive alleles contributing to liability by a factor of *c_α_* (**Fig 2D**). Fourth, we assumed that lack of robustness revealed *c_n_* additional additive alleles (**Fig 2E**). Unlike the first two models, these last two models (**Figs. 2D&E**) would allow for high heritability in families while severely confounding GWAS. Furthermore, they each increase mean and variance of the liability of the non-robust subpopulation with respect to the robust population as expected [31].

We implemented these models in our simulator (see **Fig. 3** and **Supplementary Methods** for details). Our aim was to measure the effect of changing certain parameter values, such as *n*, *φ*, *c_ρ_* and *c_n_* on expected observable values, such as the rate of concordance among monozygotic twins (*λ_MZ_*), the rate of concordance among dizygotic twins (*λ_DZ_*), and the odds ratio (*OR*) of a locus found in a typical GWAS, while holding noise (ε) constant. Our first result confirms intuition: in a mixed population, consisting of both robust (*c_α_* = 1; *c_n_* = 1) and non-robust (*c_α_* > 1; *c_n_* > 1) individuals, those crossing the clinical threshold under models D or E will be disproportionately from the non-robust subgroup. As *c_α_* or *c_n_* increase, total liability (*y*) increases. This is true for both the non-robust subpopulation *and* the entire population, which is why we use a percentile (φ) rather than an absolute threshold to determine affected/unaffected status.

**Figure 3:**
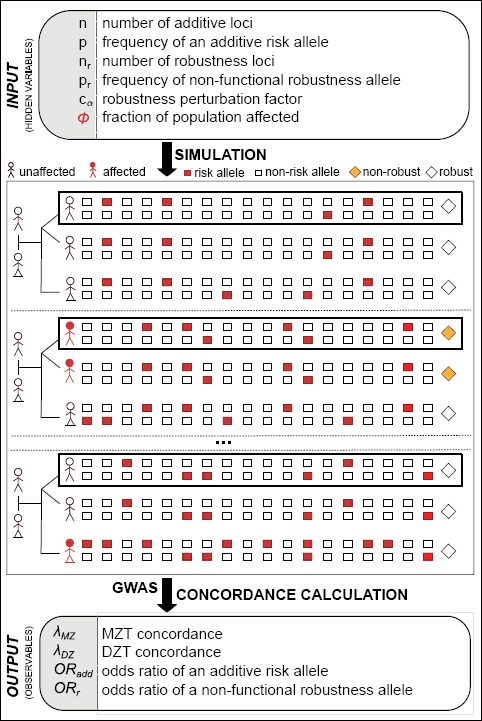
**Simulator schematic.** Using given *INPUT* parameters, we performed a simulation to generate a set of families consisting of two parents, a primary child (in bold box), and both a monozygotic and a dizygotic twin for each primary child. Each individual was assigned two alleles at random (either a risk allele (red), with probability *p*, or a non-risk allele, with probability *1-p*) for each of the *n* loci simulated. Each individual was also assigned a robustness status in the following way: if either of the two alleles at any of the *n_r_* ‘robustness’ loci were nonfunctional, each with probability *p_r_*, the individual was non-robust (indicated by gold diamond). Trait threshold is determined such that a fraction *φ* of primary children is affected. We then performed twin concordance calculations (using the simulated twins) and GWAS (using only primary individuals) to produce a set of *OUTPUT* values that we compared to empirically observed values.

**A two-state (robust/non-robust) model fits observables of real populations better than a purely additive model:** We find that a population in which just 1% of individuals are non-robust can easily produce the range of empirically determined heritabilities of binary threshold traits [**Fig. 1B inset**, *S_m_(h^2^bin)*], reported twin concordances, and broad-sense heritabilities of several complex diseases (**Fig. 1C, Table S1**). This is due to the fact that as *c_α_* or *c_n_* increase and □ remains constant, the fraction of liability that is determined by genetics (*H*^2^) increases for the non-robust subpopulation and, by extension, for the entire population.

**Controlling for robustness status adds power to GWAS:** Regardless of whether robustness is modeled as hiding cryptic variation or reducing penetrance of variants, if the goal is to find additive risk-loci (*i.e*. which may be good therapeutic targets), it is best to use *only* robust individuals for both affected and unaffected groups (**Fig. 4B**). Of course, controlling for a hidden robustness state, which modifies the effect of many alleles, returns the experiment to the situation for which GWAS was designed -- a purely additive model. It is less intuitive why using only robust (rather than all non-robust) individuals improves our ability to detect additive risk alleles in GWAS, which is a result we explore below.

**Figure 4:**
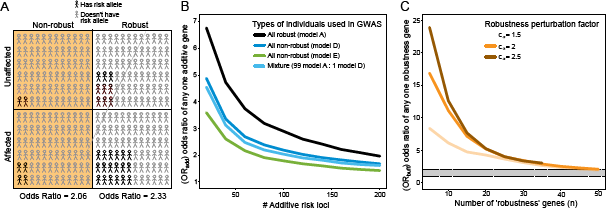
**Effect size of an additive risk locus is determined by both the total number of additive risk loci and the presence/absence of ‘gene x genome’ epistasis.** (**A**) In a robust population, where genetic liability is fully determined by the number of additive alleles, both affected and unaffected individuals will have more risk alleles than individuals of a non-robust population. Therefore the odds ratio (*OR_add_*) for any given additive allele in the robust population (2.33) will be greater than in non-robust populations (2.06), even if risk allele ratios are the same (2). (**B**) Within a population consisting of 99% robust (model A) and 1% non-robust (model D or E), the effect size of an additive allele is highest when GWAS is performed in a subpopulation consisting of only robust individuals (black line) and lowest when GWAS is performed in a subpopulation sampled without regard to robustness state (green line), with subpopulations consisting of only non-robust individuals of either type D or E performing at intermediate levels (blue lines). Input parameter values for the simulations were as follows: number of additive risk loci 20 ≤ (*n*) ≤ 200; frequency of additive risk allele (*p*) = 0.02; frequency of non-robust state (*p_r_*) = 0.01; robustness perturbation factor (*c*_α_) = 2. (**C**) Increasing the number of genes in which perturbation causes a non-robust state decreases the effect size of any one such ‘robustness’ gene to levels comparable to that most commonly found for additive risk alleles in which the risk allele is the minor allele (1 < *OR_buff_* < 2, indicated by grey horizontal bar). Input parameter values for the simulations were as follows: number of additive risk-loci (*n*) = 100; frequency of additive risk-allele (*p*) = 0.02; frequency of non-robust state (*p_r_*) = 0.01; robustness perturbation factor 1 ≤ *c_α_* ≤ 3.

In the case of cryptic variation, robust individuals have fewer available additive loci than non-robust individuals, so there are fewer ways to cross the threshold number of risk-alleles; the risk alleles are concentrated at the non-cryptic loci. Alternatively, affected robust individuals may carry additive risk alleles of larger effect size, *i.e*. a different type of risk alleles than non-robust affected individuals. Here we assume that non-robust and robust individuals carry the same additive alleles.

In the case of increased penetrance, the explanation is similar to that given for why common risk alleles are more easily found than rare alleles *that increase risk by the same amount* [15]: effect size is a function not only of the case:control ratio of allele frequency, but also of the *magnitude* of the frequency of the alleles in affected and unaffected individuals (**Fig. 4A**). If all affected and unaffected individuals are robust (as compared to non-robust), more risk alleles are required to cross the threshold (for affected individuals) and more risk alleles are allowed for individuals *not* crossing the threshold (for unaffected individuals). Therefore, while the ratio of allele frequencies between affected and unaffected individuals is the same for both all robust and all non-robust groups, the magnitude of the allele frequencies is different, making those loci easier to find in robust populations.

**Robustness status may be genetically-determined, but difficult to pinpoint:** We stated earlier that robustness status need not be genetically determined; however, we wish to explore reasonable scenarios in which it is.

Were robustness status encoded by a single genetic factor, this factor would be readily discernible, either by GWAS or linkage analysis because, under our model, affected individuals in mixed populations are disproportionately non-robust. However, robustness status is highly unlikely to be encoded by a single gene. Model organism studies suggest that a significant fraction of total number of genes – possibly up to 10% – can affect robustness [30,46,48].

To model the scenario of multiple ‘robustness’ genes, we use a simple yet plausible house-of-cards model of robustness, in which all *n_r_* ‘robustness’ loci must be functional for an individual to be robust. We observe that the odds ratio for any one ‘robustness’ locus (*OR_r_*) decreases to within the range often seen in GWAS as *n_r_ > 5* (**Fig. 4C**). In short, since there many possible ‘robustness’ loci, specific ones will be hard to find in GWAS.

With this result in mind, it may appear that models D and E are similar to the model that includes an additional additive factor of large effect (model B) in that affected individuals are disproportionately either non-robust (models D&E) or carriers of a large-effect alleles (model B) – particularly since multiple factors of large additive effect that result in indistinguishable phenotypes could conceivably work in a house-of-cards model. However, we argue that models D and E explain the marginal penetrance of some seemingly large effect factors and concomitant lack of Mendelian inheritance in pedigrees [59] as well as the presence of the multitude of other lesser risk-associated loci much more readily than model B.

## Discussion

We conclude that (i) a model that includes a house-of-cards robustness state explains the observed data for several complex diseases better than one without, (ii) when looking for loci with additive risk alleles it is best to use all robust individuals for both cases and controls, and (iii) ‘robustness’ loci are unlikely to be identified in GWAS.

In any given family, a faulty ‘robustness’ allele passed on from parent to child will act as a large-effect risk allele with Mendelian inheritance. This explains both high concordance among relatives and missing heritability in GWAS.

An obvious challenge is how to identify robust and non-robust humans. There are multiple plausible approaches that haven’t been fully explored.

In humans, one may identify ‘robustness’ genes by comparing individuals with high comorbidity of complex diseases to those with none; however, to our knowledge, this has yet to be done. Another currently available approach would be to pool individuals that are affected by distinct complex diseases and compare these to their pooled unaffected controls. As we expect non-robust individuals to be overrepresented among affected individuals for any complex disease, this GWAS approach may identify ‘robustness’ loci because the frequency of perturbed ‘robustness’ loci is increased. Indeed, this approach has shown promise for neurological disorders [60]. Similarly, the distinct diseases autism spectrum disorder and schizophrenia converge on chromatin remodeling as a general pathway that is disrupted in affected individuals [61–65]. We suggest that perturbation of chromatin remodeling leads to increased penetrance of different additive risk alleles (*i.e*. at different loci), hence resulting in distinct diseases. In fact, perturbation of chromatin remodeling genes increases the penetrance of many genetic variants in worms [30].

A third approach possible with current data would be to assume that the GWAS loci associated with the highest disease risk represent ‘robustness’ genes – particularly if there is no obvious association between gene function and a specific disease -- and proceed by controlling for these additive risk allele, recalling that statistical additivity is often an emergent property of underlying epistatic interactions [26,32,66].

Even without knowing the etiology of robustness status, it may be possible separate individuals into robust and non-robust categories. A possible proxy for robustness could be the level of genome-wide heterozygosity [67–70]. For many traits in many organisms F_1_ hybrids show less phenotypic variance and indeed hybrid vigor compared to their parental inbred lines P_A_ and P_B_ [71–75]. Plant breeders have been using this simple principle for almost a century [76]. Hybrids show decreased phenotypic variance and increased vigor despite the fact that all three populations (F_1_, P_A_ and P_B_) are isogenic and therefore all phenotypic variation *within* each population should be environmental in origin. Because phenotypic variation due to environment is apparently reduced by heterozygosity, the *fraction* of phenotypic variation due to genetics should be greater in increasingly heterozygous populations.

Some have suggested that levels of somatic genetic variation may be a read-out of robustness and mutation penetrance with higher levels of somatic variation indicating lower robustness [31,77,78]. Single-cell sequencing and phenotyping may offer insights into an individual’s robustness with greater cell-to-cell variation indicating lower robustness and higher mutation penetrance [79,80]. Non-robust individuals may also be identified as outliers in expression covariance patterns. Similar to using genotype data to elucidate population structure [81], expression data could be used to find subpopulations with different patterns of covariance between gene expression levels [82].

Robustness status may also be affected by environmental factors. Although a single environmental risk-factor should be readily identified through epidemiological studies, there could be a time lag between the environmental insult and the disease or a house-of-cards mechanism for multiple environmental insults that make them difficult to pinpoint.

Finally, it should be noted that compared to the vast resources committed to GWAS, exploring these potential markers in model organism studies seems worthwhile.

Are there indeed robust and non-robust individuals as we argue? Population genetics theory suggests that it is evolutionarily advantageous to maintain a balance of robust and non-robust individuals within a population, mainly due to the fact that non-robust individuals supply a broader phenotypic range on which selection can act under extreme circumstances [49–55].

Medical professionals intuitively agree. Drawing on their experience, they evaluate the full “Gestalt” of a patient and predict a patient’s risk for a negative outcome with often great precision. Applying ‘robustness’ markers may turn a physician’s intuition into diagnostics. Because of the potential for increasing predictive power and identifying effective drug targets, the possibility that robustness is a major player in the etiology of complex disease should be carefully considered.

## ACKNOWLEDGMENTS

We thank Joe Felsenstein for providing insightful advice that shaped this the manuscript. We thank Mary Kuhner, Cristine Alexandre. Joshua Schraiber, and Yaniv Erlich for helpful comments and discussions, and Alex Mason for help preparing the manuscript.

